# Dissecting Mutational Allosteric Effects in Alkaline Phosphatases Associated with Different Hypophosphatasia Phenotypes: An Integrative Computational Investigation

**DOI:** 10.1101/2022.01.24.477621

**Authors:** Fei Xiao, Ziyun Zhou, Xingyu Song, Mi Gan, Jie Long, Gennady M Verkhivker, Guang Hu

## Abstract

Hypophosphatasia (HPP) is a rare inherited disorder characterized by defective bone mineralization, is highly variable in its clinical phenotype. The disease occurs due to various loss-of-function mutations in *ALPL*, the gene encoding tissue-nonspecific alkaline phosphatase (TNSALP). In this work, a data-driven and biophysics-based approach for large-scale analysis of *ALPL* mutations – from nonpathogenic to severe HPPs is proposed. Allosteric molecular signatures of *ALPL* mutations were determined using an integrated pipeline of synergistic approaches including sequence and energetic-based analysis, structural topology network modeling and elastic network models. Statistical analysis of molecular features computed for the *ALPL* mutations showed a significant difference between the control and mild and severe HPP phenotypes, and the developed machine learning model suggested that the topological network parameters could serve as is a robust indicator for severe mutations. Molecular dynamics simulations coupled with protein structure network was employed to analyze the effect of single-residue variation on conformational dynamics of TNSALP dimers, and we found that severe disease-associated mutations have a significantly greater effect on allosteric communications. The results of this study suggested that *ALPL* mutations associated with mild and severe HPPs tend to have different effects on protein stability on local scale and long-range communications caused by network rewiring. By linking disease phenotypes with allosteric molecular signatures, the proposed integrative computational approach elucidates the complex sequence-structure-allostery relationships of *ALPL* mutations and dissects the role of allosteric effects in the pathogenesis of HPPs.

**Author Summary:** The understanding of mutational genotype- disease phenotype relationship is a fundamental step for enabling precision medicine. It remains a challenging task to assess the molecular principle of the genotype- phenotype relationship. By focusing on Hypophosphatasia, a rare inherited disorder, as an example, we performed comprehensive analysis of single-amino-acid mutations in the encoded protein of Tissue Nonspecific Alkaline Phosphatase associated in terms of their sequence, structure, and dynamics properties. We further developed a machine learning -based method to classify different disease phenotypes, and the interpretability of the classification model was addressed by the structural-functional analysis of network topological important mutations. Our results highlighted the allosteric propensity of severe mutations, and show the potential allosteric principle of genotype- phenotype relationship.

## Introduction

Hypophosphatasia (HPP) is a rare autosomal dominant or recessive metabolic disorder, first described in 1948 by Rathbun,[1] which constitutes a complex, multisystemic disease.[2] The clinical presentation is highly variable, depending on the type of mutation and the inheritance mechanism. From a prenatal lethal form with no skeletal mineralization to a mild form with late adult onset, HPP can be classified into six subtypes, including perinatal, infantile, prenatal benign, childhood, adult and odonto.[3] In general, perinatal and infantile subtypes represent severe forms of HPP, while childhood, adult, odonto, and prenatal benign constitute mild phenotypes.[4] HPP is caused by loss-of-function mutations in the *ALPL* gene encoding the Tissue Nonspecific Alkaline Phosphatase (TNSALP).[5] TNSALP is a membrane-bound metalloenzyme, whose activity is reduced by various mutations in the *ALPL* gene, leading to increased inorganic pyrophosphate, which in turn cause different HPP phenotypes.[6] As an enzyme replacement therapy, ENB-0040 is a bone-targeted, recombinant human TNSALP that prevents the manifestations of HPP.[7] However, to date, there is no established treatment for HPPs,[8] due to the little knowledge about the relationship between mutations in the gene responsible for HPPs and phenotypic variability.

The *ALPL* gene is localized on chromosome 1p36.1–34 and consists of 12 exons distributed over 50 kb.[9, 10] *ALPL* mutation detection is important for recurrence risk assessment and prenatal diagnosis.[11] To date, more than 400 *ALPL* gene mutations have been reported worldwide and about 80% of these mutations are missense. Although HPP is caused by homozygous, heterozygous, or compound heterozygous *ALPL* mutations,[12] most of these mutations cause change in a single amino acid in TNSALP. Mutations in *ALPL* would reduce the enzyme activity of TNSALP mutant proteins to a varying degree, with residual activities often exhibiting different enzymatic properties from wide-type TNSALP. By measuring the effects of amino acid changes on TNSALP dimer stability, the relationship between *ALPL* missense variants and TNSALP enzymatic activity can be predicted.[13] Despite recent progress, the correlation between these mutations and the six HPP subtypes is not well established, and thus the molecular mechanisms underlying the genotypic-phenotypic relationships of HPP remain unclear.[14] A more detailed molecular characterization of *ALPL* mutations can help to establish the etiology, pathogenesis, and onset of HPP. Moreover, understanding molecular signatures of pathogenic mutations may provide a better strategy for inhibiting TNSALP and tools to govern rationale molecular targeted strategy.[15] TNSALP is a homodimeric protein that contains several known domains, including five principal functional domains: a catalytic site, calcium-binding site, crown domain, homodimer interface, and N-terminal alpha helix.[12] The assignment of various mutations to the five functional domains of the TNSALP structure model has contributed to the current understanding of the genotypic and phenotypic interrelationship of HPP.[16] For example, most of the severe missense mutations were localized in crucial domains, such as the active site, the vicinity of the active site, and the homodimer interface. The structural importance of the crown domain has also been highlighted, with respect to the catalytic function of TNSALP.[17] In addition, there is structural evidence to support the concept that these crucial domains are also involved in allosteric properties of TNSALP, since the catalytic activity depends on its homodimeric configuration from which the dominant negative effect of some loss-of-function ALPL mutations is derived. Although mammalian alkaline phosphatases have been known as allosteric enzymes for many years,[18] the dynamics-driven allosteric signaling pathways is yet to be investigated at an atomistic level. Therefore, the elucidation of the conformational dynamics and allosteric signatures of TNSALP caused by *ALPL* mutations would provide greater insight into genotype-phenotype relationship in HPP.[19, 20] In order to predict catalytic efficiency[21] and potential pathogenicity of mutations,[22, 23] different levels of computational methods, from sequence analysis to structural and dynamics analyses have been developed. From a molecular evolution perspective, sequence-based methods have shown that pathogenic mutations always involve conserved amino acids.[24] Furthermore, it has been pointed out that disease-causing mutations frequently involve a drastic change in amino acid physicochemical properties, such as charge, hydrophobicity, and geometry, and are fewer surface exposed than polymorphic mutations.[25] Structural modeling combined with network theory has been widely exploited in studying protein topology and dynamics.[26, 27] The description of protein structure as a structural network provides an efficient tool to dissect complex protein functions.[28] Based on protein structural network model, different topological parameters and algorithms have been developed to describe structural changes associated with mutations.[29, 30] Conformational dynamics would provide more insight into the physical basis of the effect of missense variants.[31] Elastic network models (ENMs) have long been established in the analysis of protein dynamics in the context of normal model analysis,[32, 33] and have now become an extremely useful approach to predict the effects of single-point mutations on protein stability.[34] The calculation has revealed that mutations are significantly enriched on hinge-neighboring residues in oncogenes and tumor suppressor genes.[35] By incorporating dynamics descriptors based on ENM with sequence and structure-dependent properties, the prediction accuracy of impact of variants on biological function has greatly increased.[36] Machine learning models that integrate sequence, structure, and ensemble-based features have been developed to classify mutation types.[37–39] Furthermore, biophysical simulations combined with structure-based modeling of residue interaction networks have also been used to reveal the functional role of mutation hotspots in molecular communication in some tumor suppressor proteins,[40] regulatory complexes including HSp90[41], and SARS-CoV-2 Spike Protein,[42] as well as classify *PTEN* missense variants corresponding to cancer or autism spectrum disorder.[43–45]

Our recent works have highlighted that biophysics-based and data-driven approaches, including genomic analysis, coevolution and network-based modeling provide an array of powerful tools to study disease mutations in genomic medicine and allosteric interactions.[46–49] In this work, based on this concept, we propose integrated computational approach for the large-scale analysis of disease mutations. As shown in Fig 1, we collected diverse *ALPL* single-point mutations associated with neutral, mild, and severe phenotypes from various scientific gene mutation databases; the computational pipeline consisted of three steps. First, by performing three levels of analysis, including conservation and co-evolution analysis based on sequence level, structural modeling, energetic analysis and protein structure network (PSN) modeling based on structural level, and ENM-related calculation based on dynamics level, different molecular and network signatures were attributed to three types of mutations. Second, statistical analysis including creation of a random forest mode, was performed to test which molecular signatures can serve as robust predictors for classifying the three kinds of mutations. Finally, through integration of long-range perturbation dynamics and network-based approaches, we have quantified allosteric potential of selected mutation residues. Our study characterizes the mutational landscape of *ALPL* through modeling and analysis of molecular signatures and allosteric effects of mutations,[50] providing new insights into the genotype-phenotype interrelationship in HPP.[51]

**Fig 1.**
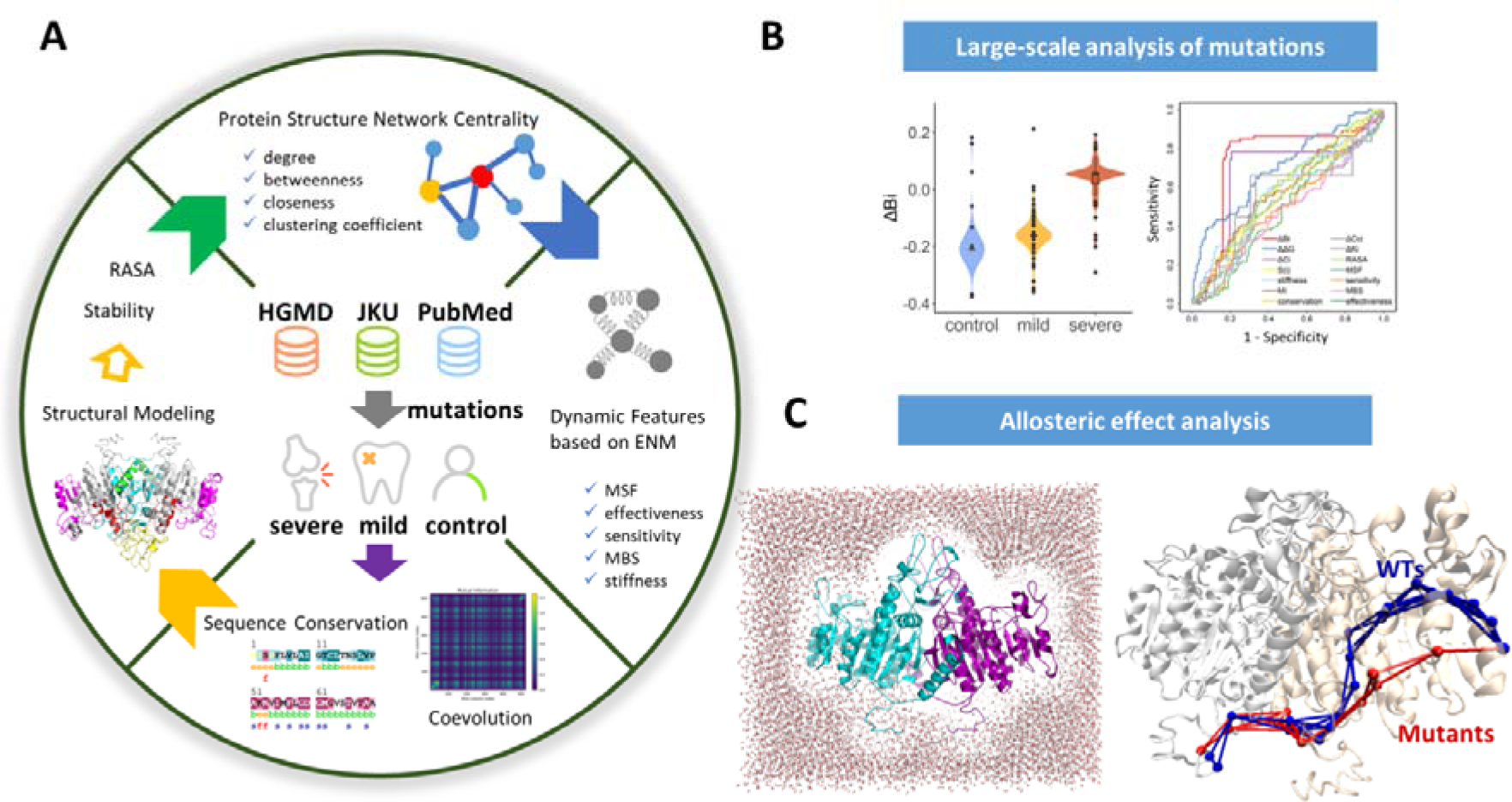
The computational workflow. (A) Beginning with the collection of *ALPL* mutations associated with different HPP phenotypes and the computational modeling of TNSALP protein structure, molecular signatures of mutational hotspots were calculated. In addition to normally used signatures, three levels of parameters for describing mutations were analyzed: conservation and co-evolution analysis at sequence level, PSN-based network matrices at structural level, ENM-related features at dynamics level. (B) The classification and prediction of pathogenicity of *ALPL* mutations based on the statistical analysis of molecular signatures and the construction of machine learning models. (C) The allosteric effect analysis of predicted mutations by single-residue perturbation, molecular dynamics and long-range pathway analysis.

## Materials and Methods

### Data Collection of ALPL Mutations

With the goal of classifying various *ALPL* variants according to different phenotypes, 242 single-point loss of function variants were selected in HPP patients with mild and severe phenotypes and the control group from the *ALPL* gene mutation database (https://alplmutationdatabase.jku.at, accessed on 11 January 2021), the Human Gene Mutation Database (http://www.hgmd.cf.ac.uk/ac/index.php), the Locus Specific Mutation Databases (http://www.hgmd.cf.ac.uk/docs/oth_mut.html), and literature in the PubMed database. The collated data set of *ALPL* mutations associated with mild and, severe HPP phenotypes and putative functionally nonsense variants, as well as their molecular signatures, are listed in S1 Table.

### Sequence Conservation and Coevolution Analyses

By using the Consurf server,[52] the conservation scale for each residue in the TNSALP protein was calculated (range from 1 through 9, where 1 denotes the node least conserved and 9 denotes the most conserved sites). The refined multiple sequence alignment (MSA) of the TNSALP protein can also be retrieved from the Consurf server, which has been subjected to Shannon information entropy and mutual information (***MI***) calculations. measures the variability of specific sites of protein sequences, which is calculated as

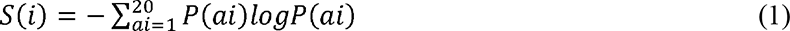

where is the probability of occurrence of amino acid type at the *i^th^* column. varies in the range 0≤ ≤3.0, a lower implies higher evolutionary conservation. Similarly, *MI* was applied as a measure of the degree of intra-molecular co-evolution between residues. The *MI* associated with the i^th^ and j^th^ sequence positions is defined as an matrix of the form

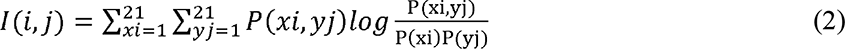

where P(xi, yj) is the joint probability of observing amino acid types *x* and *y* at the respective sequence positions, *i* and *j*; P(xi) is the marginal/singlet probability of amino acid of type *x* at the i^th^ position. I(i, j) varies in the range [0, I_max_], corresponding to fully uncorrelated and most correlated pairs of residues. The co-evolution of a mutation was measured by the average *MI* values corresponding to each residue. The calculation of and *MI* were performed by Evol.[53]

### Structural Modeling and Protein Stability Analysis

A protein homology model for human TNSALP was constructed by using the MODELLER V9.19[54] platform, using a template corresponding to the human placental alkaline phosphatase (PDB id: 1ZED[55]). The template had a sequence identity of 57% to TNSALP and an X ray crystal structure resolution of 1.57 Å. Each single *ALPL* missense variant and its mutant structures were automatically generated by FoldX,[56] and its effect on protein stability can be measured by the difference folding Gibbs free energy (ΔΔ***G*** values) between the wild type (WT) and the mutated forms of TNSALPs. The accessible surface area (ASA) is the atomic surface area of a protein that is accessible to a solvent, and the relative ASA (***RASA***) attribute is the per-residue ratio between the calculated ASA and ’standard’ ASA for a particular mutational residue. As a measure of amino acid side-chain accessibility, the RASA also serves a quantitative predictor of variants’ pathogenicity.[57] The RASA calculation of TNSALP is done through PSAIA 1.0.[58]

### Protein Structure Network Analysis

Protein structure networks for the TNSALPs were constructed and analyzed by using the NACEN R package.[59] A node in the network denotes a single amino acid residue, and edges are defined by the environment-dependent residue contact energy between two nodes.[60] Based on PSNs, some network centralities have been defined.[61] The simplest centrality measure is the degree (***K****_i_*) of a node *i* in PSNs, defined as the total number of nodes that it is directly connected to. The betweenness centrality ***B****_i_* was defined as the number of times residue *i* was included in the shortest path between each pair of residues in the protein, normalized by the total number of pairs. It is calculated by

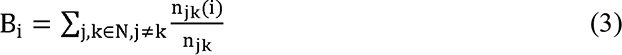

where n_jk_ is the number of shortest paths connecting *j* and *k*, while n_jk_is the number of shortest paths connecting *j* and *k* and passing through *i*. The closeness centrality ***C****_i_* for a node was calculated by the reciprocal of the average shortest path length, which can be calculated as follows:

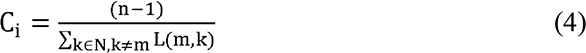

where N is the set of all modes and *n* is the number of nodes in the network. The clustering coefficient (***Cc****_i_*) measures the degree to which nodes tend to cluster together and is defined as:

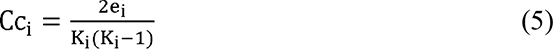

where *K_i_* is the degree of node *i* and *e_i_* is the number of connected pairs between all neighbors of *i*.

### Dynamic Features based on Elastic Network Models

In ENMs,[32, 33] each node represents a Cα atom in proteins and each edge is a spring γ for connecting two sites within a given cutoff distance r_c_. Two most commonly used ENM methods, the GNM and the ANM, are adapted in this paper. The total potential energy of the ANM and GNM systems with N nodes are expressed as

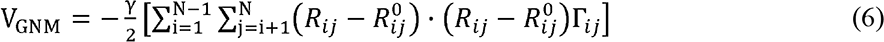

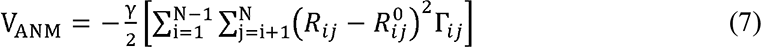

where R and R are the instantaneous and equilibrium distances between nodes i and j, andis the ijth element of the N N Kirchoff matrix Γ, which is written as

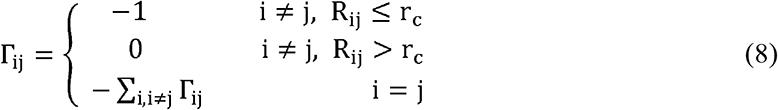

In our study, r_c_ between protein nodes are 7 Å and 13 Å for GNM and ANM, respectively. In comparison with GNM that only measure fluctuation, ANM provides additional information of motion directions of each residue. Herein, the GNM and ANM calculations were performed by ProDy.[62]

The normal modes are extracted by eigenvalue decomposition Γ U1U$. U is the orthogonal matrix whose kth column Uk is the kth mode eigenvector, and 1 is the diagonal matrix of eigenvalues, λ . Mean-square fluctuations (MSF) of a residue are given by

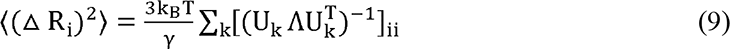

where k& and T represent Boltzmann constant and temperature, respectively. Based on ENM calculation, several other kinds of matrixes can be generated. Form Perturbation Response Scanning (PRS) matrix,[63] two dynamic features of ***effectiveness*** and ***sensitivity*** were defined as row and column averages of the matrix, respectively. The effector residues most effectively propagate signals in response to external perturbations. The sensor residues can easily sense signals and respond with dynamic changes. Directly from Kirchoff matrix, its eigenvalues can be used to ascertain how important each node is to maintain the overall mechanical connectedness of the network. This amounts to measuring how much the network Laplacian spectrum changes when the connections, or couplings, of a node with its neighbors are deleted. As such, the Mechanical Bridging Score (***MBS***) for a given mutation reflects the response ability of this residue. In the symmetrical stiffness matrix, the elements describe the effective spring constants associated with each residue pair. The ***stiffness*** for individual mutational residues is obtained by averaging all the elements in the corresponding row/column of the matrix. The detailed description of these dynamic features can be found in Bahar’s work.[36] The statistical analysis of all features was performed in the R language (https://www.r-project.org/, R Core Team, 2017), version 3.6.2. Plots were generated using R package ggplot2, version 3.3.3.[64] Statistical significance was determined by the Wilcoxon signed ranked test (P <0.01) in our analysis.

### Molecular Dynamics Simulations

The MD simulations were performed on WT and mutant TNSALPs for the conformational dynamics study. To eliminate any unfavorable contacts, energy minimization was performed. Each structure was first minimized with the steepest descent algorithm and followed by the L-BFGS algorithm without any restraint. Total minimization was carried out until convergence, where the maximum atomic force was <500 kJ/mol-nm. The minimized structures were then slowly heated from 0 to 300 K over 100 ps and equilibrated for an additional 200 ps. Later, MD simulations were further carried out for 100 ns using the GPU-enabled version of the GROMACS software package (version 5.1.4)[65] with the AMBER99SB-ILDN force field,[66] using a timestep of 2 fs. The structures were immersed in an octahedral box filled with TIP3P water molecules, imposing a minimum distance of 15 Å between the solute and the box. The non-bonded interaction potential was smoothly switched off between 10 and 12 Å, beyond which coulombic interactions were treated with the particle-mesh Ewald method.[67] The LINCS algorithm[68] was used to constrain the hydrogen containing bonds. A temperature of 300 K was maintained via the V-rescale dynamics.[69] The atomic positions were saved every 5000 steps (10 ps) for analyses. GROMACS analysis toolkit utilities were used to analyze MD trajectories produced during the last 70 ns production run to determine root mean square deviations (RMSDs), root mean square fluctuations (RMSFs) for each system.

### Allosteric Mutation Analysis

Two methods were used for the investigation of allosteric effects of mutations. The dynamic network analysis was performed by the program, MD-TASK.[70] For each protein system, the network was constructed by extracting C_α_ and C_β_ atoms of MD trajectories as nodes and the edge is created when two nodes are within 6.5 Å. Then, the allosteric communication pathways were calculated by connecting mutational sites and allosteric sites with the shortest edges, using the Floyd-Warshall algorithm.[71] AlloSigMA server[72, 73] was used to evaluate the allosteric effects of each mutation based on the Structure-Based Statistical Mechanical Model of Allostery. This statistical mechanical model estimates the allosteric free energy difference Δg of each residue used by the allosteric communication under mutation events to perform the allosteric mutation analysis in TNSALP.

## Results and Discussion

### Sequence and Structural Landscape of ALPL mutations

To clarify the sequential and structural characteristics of genotype and phenotype, we first divided all mutations into three categories according to the severity of HPP: the mild form, severe form, and control group. Here, 185 missense mutations related to 142 unique amino acids were found in the *ALPL* gene, of which 111 mutations related to 100 residues were found to cause the mild form of HPP, while 74 mutations related to 63 residues were found to cause the severe form of HPP (Fig 2A). Moreover, 57 mutations related to 55 residue sites were found in asymptomatic people and, therefore, served as the control group. It should be noted that a residue may be related to different types of mutations (S1 Fig), while the Venn Diagram showed overlap of mutated residues of all three groups (Fig 2A). The primary sequence analysis showed that there are 261 conserved residues (conservation score>5, see S2 Fig) among the full length of 524 residues of the TNSALP monomer. In order to characterize the relationship between conservation and phenotype, we calculated the proportion of conserved residues in each group, and results are shown in Fig 2B. Among the mutation sites that cause severe phenotypes, more than 71.0% of the sites are highly conserved, while in the mild group, this proportion is 65.0%, and in the control group, it is much lower, at only 27.3%.

**Fig 2.**
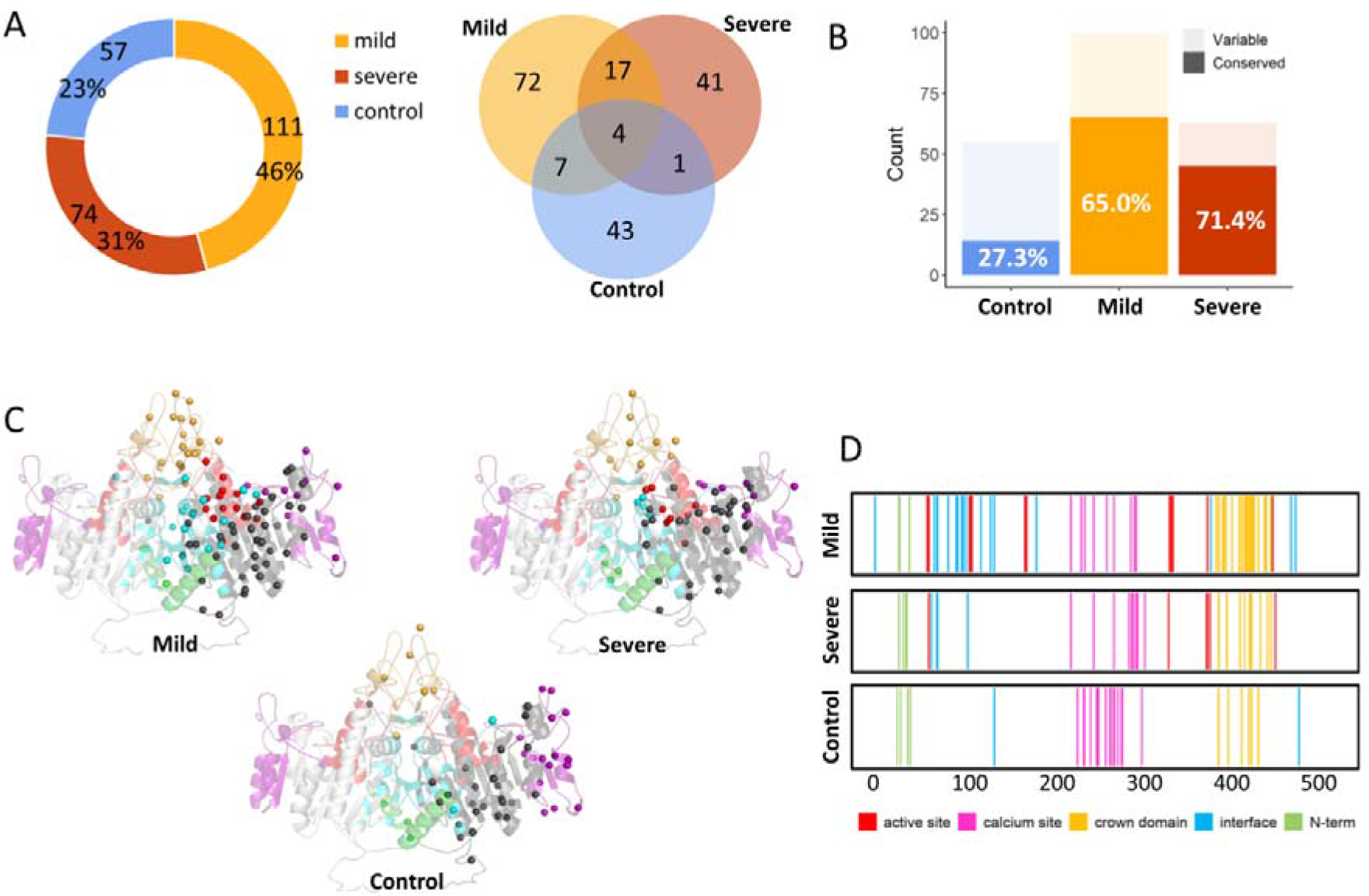
Sequence and structural analysis of *ALPL* mutations in the TNSALP protein. (A) The composition and statistics of *ALPL* mutations of based in the three categories: mild HPP, severe HPP, and control group. (B) The conservation distribution of mutations related to different phenotypes. (C) The structure of the TNSALP protein, and the distribution of mutation sites in the active site (red), calcium site (purple), crown domain (brown), dimeric interface (blue), and N-terminal domain (green). Missense mutations in the three groups are shown as colored spheres based on the coloring scheme of the domains to which they belong. (D) Distribution of different clinical phenotypic mutations across the length of the TNSALP protein.

It is worth noting that half of these highly conserved sites in the control group are from the intersection of the mild group and the severe group. Thus, we postulated a possible trend that mutations occurring at highly conserved sites are more likely to cause HPP disease.

The 3D structure of TNSALP was then computationally modeled by using the crystal structure of human placental alkaline phosphatase as the template (S3 Fig). By mapping all the mutations onto the modeling TNSALP structure, we identified the spatial distribution of the three kinds of mutations (Fig 2C and 2D). It was found that 104 out of 185 (66%) pathogenic mutations were located at these five functional domains; in particular, 17% were at the crown domain, while only 28 of 57 (49%) mutations in the control group were distributed in this domain. In the mild group, 12 mutations were located at the active center, 21 at the interface, 23 at the crown domain, 9 at the Ca^2+^ binding site, and 3 at the N-terminal. The distribution of mutations in the severe group is as follows: 7 mutations at the active center, 4 at the interface, 9 at the crown domain, 12 at the Ca^2+^ binding site, and 4 in N-terminal (Fig 2D). On the other hand, in the control group, 16 mutations were located at the Ca^2+^ binding site, 2 at the interface, 6 at the crown domain and 4 at the N-terminal. The major difference in the spatial distribution between pathogenic and control groups was that the control group had no mutations at the active site, thereby suggesting the role of active sites in HPP pathogenicity.

### Structural Environment and Evolutionary Properties of ALPL Mutations

The overall sequence and structural landscape of *ALPL* mutations suggest that evolutionary conservation and spatial distribution of mutation sites in the TNSALP protein may be linked to different functional roles in disease phenotypes. Here, we performed the comparative RASA and evolutionary calculation of TNSALP for the large-scale analysis of *ALPL* mutations. These physicochemical features could impart a particular structural environment and evolutionary properties to mutational hotspot sites. According to the RASA (Fig 3A), residues were classified into three classes, i.e., buried areas (RASA<5%), semi-exposed areas (between 5% and 30%), and fully exposed areas (RASA>30%). Although different types of mutations were distributed across all functional domains, disease-causing variants are more likely to occur at the buried areas (122 mutations, 65.9%). By contrast, nonsense variants are more likely to occur at the surface (32 mutations, 56.4%). We systematically characterized the evolutionary landscape of *ALPL* mutations; the distribution of the three types of mutations on and ***MI*** profiles are shown in Fig 3B and C, respectively. Regarding conservation, 178 of the 524 amino acid residues in the protein sequence are relatively conserved (<1.0). In general, disease-causing variants are more likely to occur at the conserved sites (99 mutations out of 185 pathogenic mutations with <1.0, 53.5%), while the nonsense variants are more likely to occur at the less conservative sites (44 mutations out of 57 neural mutations with >1.0, 77.2%). We observed a significant positive correlation between co-evolution and entropy (S4A Fig), and the distributions of and ***MI*** scores for mutations were also very similar. By using ***MI*** = 0.2 as a cutoff, the low values of amino acids correspond to residues that were highly co-evolutionary. For pathogenic mutations, 64.9% (48 out of 74) severe mutations and 55.9% (62 out of 111) mild mutations occurred at highly co-evolutionary residues, whereas for the control mutations, only 28% belonged to co-evolutionary residues.

**Fig 3.**
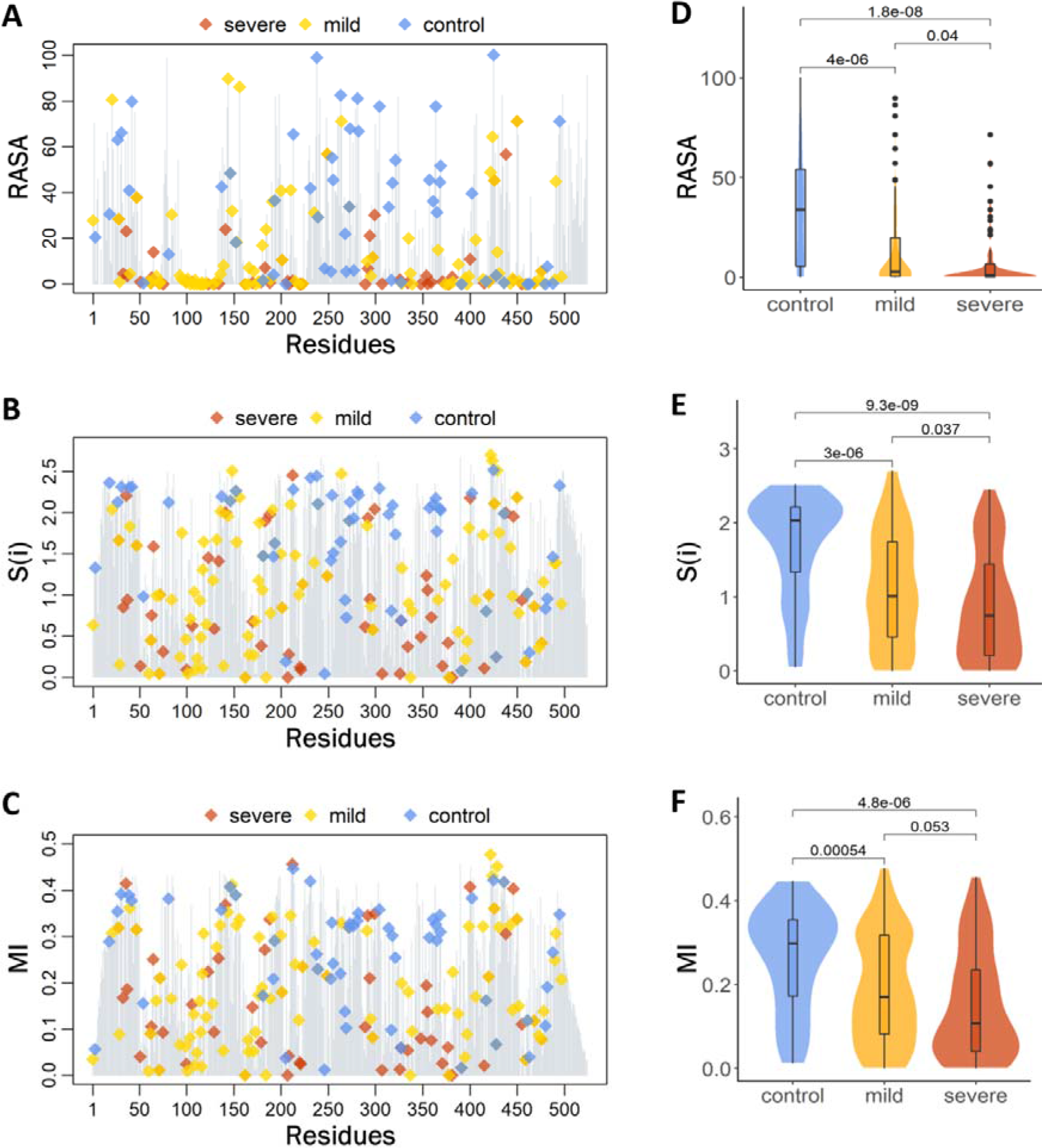
The (A) relative solvent accessible area (RASA), (B) entropy ( ) and (C) co-evolutionary (*MI*) profiles for each residue in TNSALP; mild, severe, and control mutations are highlighted as yellow, red, and blue diamonds, respectively. The comparison of (D) the RASA, (E) and (F) *MI*, between the control group mutations and mild and severe mutations. Statistical significance was determined by the Wilcoxon signed-ranked test, with *P* values 0.01.

The statistical analysis of RASA, and ***MI*** for control, mild, and severe mutations showed statistical significance between the control and mild, and control and severe mutations; however, there was no statistical significance between the mild and severe mutations (Fig 3D-F). Accordingly, structural environment and evolutionary studies have found that the HPP-causing *ALPL* mutations, especially severe mutations, are located in buried, conserved and co-evolutionary residues with small RASA, and ***MI*** scores.

### Energetic Analysis Reveals the Effect of Severe and Mild Mutations on TNSALP Stability

To explore the thermodynamic determinants of mutational hotspots and establish a robust baseline for ranking and evaluating functional significance of mutational hotspots, FoldX[56] was used for the prediction of free energy changes in single point mutations. In this method, a mutation is predicted when there is a destabilization effect positive value of Gibbs free energy changes (ΔΔG), while the respective mutation is a stabilized protein with a negative value of ΔΔG. As shown in Fig 4(A), we found that most of the disease mutations are associated with decrease in binding affinity (ΔΔG,0), reducing the protein stability. Several mutations showed significant peaks corresponding to the severe phenotype, which were located in different domains, such as G220R, G326R, G326V, G221R, G221V, D378H, G129, G334D, C201Y, R223W, and G63R. We also noticed that the largest destabilization effect upon substitution was found for mutations in the mild group, namely G339R, P108L, A468V, G491R, C139Y, A111T, R428Q, A114T, and G249V. On the other hand, the mean value of ΔΔG for most mutations in the control group, such as L365M, K27S, V314E, S368Q, R272C, R246S, L18F, and Y39N was < 1.0 kcal/mol. Furthermore, as shown in Fig 4B, we found that the predicted folding free energy change ΔΔG of mutations which cause severe phenotype (mean ΔΔG was 5.82 kcal/mol) was obviously higher than that in the mild group (mean ΔΔG was 2.05 kcal/mol, *P*=2.4e-06), and much higher than that in the control group (mean ΔΔG was 0.43 kcal/mol, *P*=1.5e-11). Therefore, most of the disease-causing mutations are associated with decrease in binding affinity, and this trend varies in different phenotypes.

**Fig 4.**
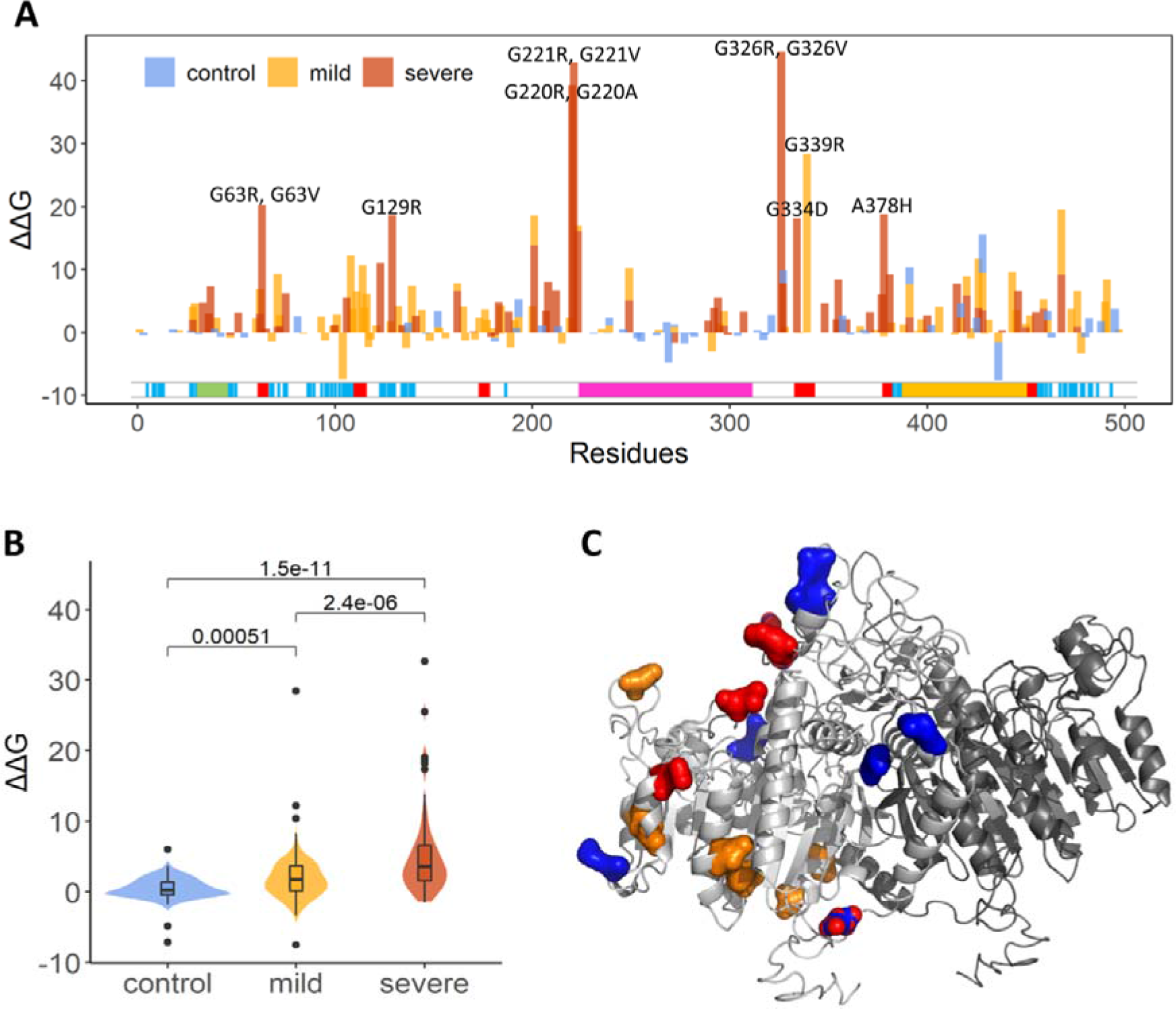
(A) The free energy changes induced by single point mutations. The profiles of mild, severe, and control mutations are shown as yellow, red and blue bars, respectively. (B) The statistical difference between predicted folding free energy change of *ALPL* single residue variations in the control, mild, severe groups was measured by Wilcoxon. test, with *P* values 0.01. (e) The structural distribution of disease-causing mutations with low ΔΔG.

In addition, ΔΔG provides a benchmark or metric that can be used to compare with physicochemical properties for large-scale analysis of other mutations. We have compared the ΔΔG with the RASA, and ***MI***, and weak correlations were found among them (S4B-D Fig), with negative coefficients of 0.285, 0.350 and 0.321, respectively. These low coefficients imply that the three properties provide additional measures of ΔΔG for describing *ALPL* mutations, namely pathogenic mutations tend to have a high ΔΔG value and low RASA, *S*_(*i*)_, and ***MI*** values. For example, pathogenic mutations, including G220R, G221R, G326R, G129R, P108L, R223W, and G63R (severe mutations), and G339R and A468V (mild mutations) showed obvious energy change but with low RASA, *S*_(*i*)_, and ***MI*** scores. By contrast, several nonsense mutations, such as L18F, R272C, H281S, E146K, and T273M, were predicted to have small ΔΔG but with high RASA, *S*_(*i*)_, and ***MI*** scores. Based on the above analysis, we concluded that most of the “severe” and “mild” mutations were predicted to have large ΔΔG values and differed in terms of evolutionary conservation and physicochemical properties.

However, some mutations were not consistent with the above results. In the severe group, mutations including R272L, N47I, L299P, R272H, H438L, and R450C, the ΔΔG values of many mutations were < 1.0 kcal/mol, while the RASA score was > 30%, > 1.0, and MI > 0.2. Mild group mutations, such as R272H, R450H, V424A, K264R, E21K, V424M, R184Q, E84V, N47I, K422R, and T148I showed a slight energy change (ΔΔG < 1.0 kcal/mol) but displayed high co-evolutionary conservation (>1.0, MI >0.2) and RASA score (>30%). By contrast, some nonsense mutations such as I269V, H482N, F327C, A488S, T277A, T255I, I317L, and D239Y, were predicted to have a large energy change (ΔΔG > 1.0 kcal/mol) with low co-evolutionary conservation scores (<1.0, MI <0.2) and RASA score (<30%), Structurally these mutations are distributed at different domains and not clustered together (Fig 4C). These results highlighted potential limitations of using only the folding free energy (ΔΔG) changes to predict pathogenic mutations, thus suggesting that molecular disease mechanisms may be determined not only by local energetic changes but through impairment of long-range allosteric interactions and alterations in the global interaction networks. In addition, the random structural distribution of disease-causing mutations calls for a structural systems biology approach for predicting the effect of different types of mutations, toward the understanding of their genotype-phenotype relationships.

### Network Modeling of *ALPL* Mutations

Network theory has become a widely used method to quantify protein structures and functions in terms of their topological connectivity.[74] In our analysis, we used the difference between network centrality measures of mutant and WT values to describe the topological change of TNSALP caused by *ALPL* mutations. For each amino acid substitution, four common network parameters including degree (K_i_), betweenness (B_i_), closeness (C_i_), and clustering coefficient (C_Ci_) were calculated. The computational results of ΔK, ΔB, ΔC, and ΔC_C_ associated with each mutation in the mild, severe, and control groups, are illustrated in Fig 5 and 6.

**Fig 5.**
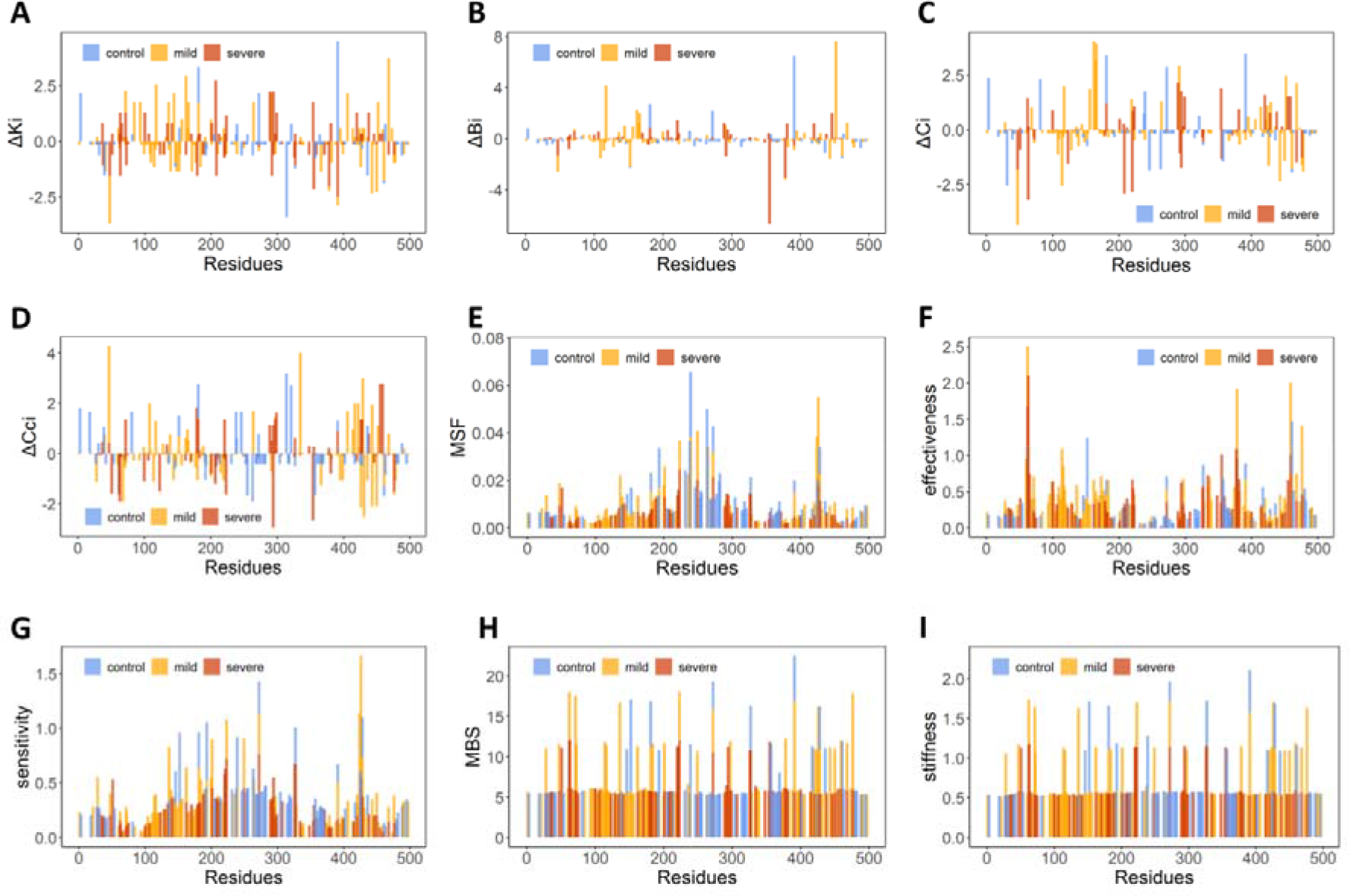
Network topological and dynamic analysis of mutations in *ALPL*. (A)ΔK, (B) ΔB, (C) ΔC, (D) ΔCc, (E) MSF, (F) Effectiveness, (G) Sensitivity, (H) MBS, and (I) stiffness profiles for the three types of mutations. Control, mild and severe mutations are represented as blue, orange, and red bars, respectively.

**Fig 6.**
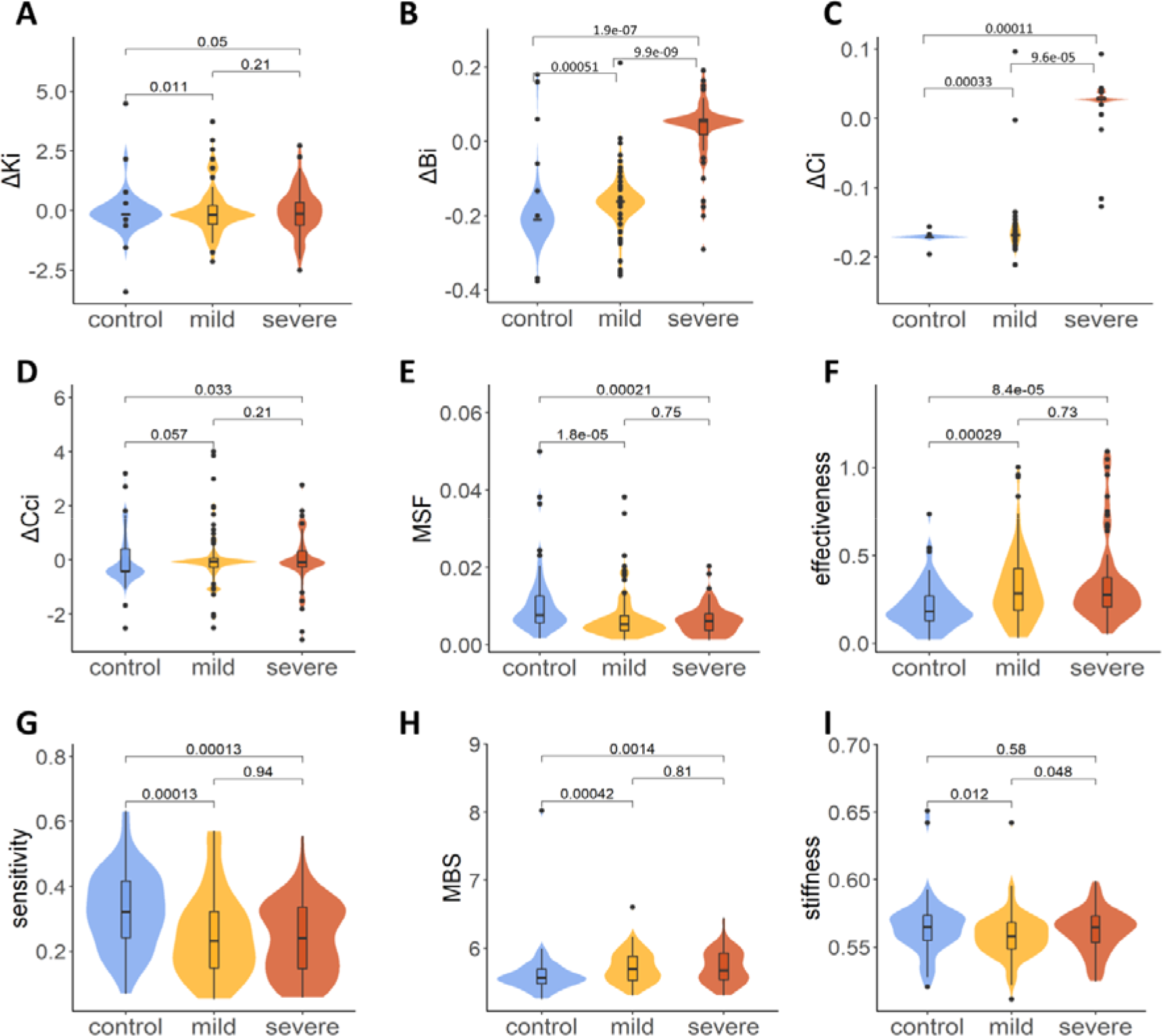
Comparison of the change of four network centralities and five dynamics-based parameters, including (A)ΔK, (B) ΔB, (C) ΔC, (D) ΔCc, (E) MSF, (F) Effectiveness, (G) Sensitivity, (H) MBS, and (I) stiffness, among mutations in the control, mild, and severe groups by Wilcoxon signed ranked test.

The profiles in Fig 5 (A-D) depict that most of the mutations show a wide range of changes in the values of K_i_, C_i_, and C_Ci_. In general, some common peaks were found in ΔK, ΔB, and ΔC profiles, such as the severe mutation R391H and three mild mutations G162S, T166I, and R272C. Detailed analysis of the ΔK profile showed that its peaks corresponded to pathogenic mutations, including severe mutations R391C, Q207P, D294A, L289F and mild mutations A468V, G162S, Y117C, G144E, E291K, and D406G. These significant peaks aligned precisely with functional centers, including hotspot positions involved in the crown domain, the homodimer interface, and the Ca^2+^ binding site, thereby suggesting their latent functional roles. In comparison, smaller number of variations were observed in the ΔB profile (Fig 5B), but much sharper peaks for severe mutations F355I, D378H and A446T and mild mutations E452K, Y117C, G162S, R152C, and T166I were found. Functional domain analysis revealed that E452 and D378, R391, and A446, R272 are involved in the active site, the crown domain, and the Ca^2+^ binding site, respectively. Eleven peaks were found in both the ΔC and ΔCc profiles. In the ΔC profile (Fig 5C), eight corresponded to pathogenic mutations, including G162S, T166I, G63R, T165I, L208F, E291K, G220R, and N47I, and only one (E291) was located at the functional domain. In the ΔCc profile (Fig 5D), the other set of pathogenic mutations, peaks corresponded to R335T, N47I, E429K, D294Y, G455D, V459F, E354D and V431A, including many functional sites. Among these, R335 and G455 were located at the active site, D294 and Y263 at the Ca^2+^ binding site, and E429 and V432 at the crown domain.

On the statistical comparison of the topological differences between the three groups of mutations, it was observed that four network parameters were decreased for most groups of mutations (with mean values <0), only ***B****_i_*and ***C****_i_* were increased for the severe mutations. In addition, both ΔB and ΔC showed significant differences between any two groups of mutations (Fig 6). As shown in Fig 6 (B), the mean values of ΔB for the severe, mild, and control groups are 0.05, 0.16 and 0.21, respectively, with *P*=1.889e-07, 9.906e-09, and 0.0005129, respectively, by the Wilcoxon signed rank test between the severe and mild, severe and control, and mild and control groups.

In contrast, as shown in Fig 6 (C), the mean values of ΔC for the severe, mild, and control groups are 0.03, 0.17 and 0.17, with *P* = 9.646e-05, 0.0001094, and 0.0003274, respectively, by the Wilcoxon signed rank test, between the severe and mild, severe and control, and mild and control groups. Through this network centrality analysis, we found that pathogenic *ALPL* mutations showed significantly higher variation in ΔB and ΔC compared with the control group. As two network global topological measures, the significant change in betweenness and closeness suggests that pathogenic mutations (especially severe mutations) may affect the overall connectively of the interacting network rather than the local geometry. In particular, ΔB showed the best discrimination, as has been suggested, and could serve as a potential in-silico marker for pathogenic NaV1.7 mutations.[75] Remarkably, the increase of betweenness and closeness caused by severe mutations may suggest a structural and functional change of the protein, by tightening the correlated dynamical network and making the severe state as a more allosteric molecular machine.

### Elastic Network Modeling Highlights Mutational Signatures of Conformational Flexibility

In addition to sequence conservation and structural information, structural dynamics has also been shown to be another determinant of the functional impact of missense variants. A series of ENM-based dynamics descriptors, including GNM-based MSF, PRS-based effectiveness and sensitivities, and ANM-based mechanical bridging score (MBS) and stiffness, have been introduced to enhance the predictive ability of disease-related mutations[36]. Here, we employed these dynamic features to systematically characterize different types of *ALPL* mutational sites.

First, mapping the three types of mutations onto the dynamic profiles based on the TNSALP protein, provided primary insight into the interpretation of the functional impact of variants in the light of the intrinsic dynamics of the mutational site (Fig 5 E-I). As shown in the MSF profile (Fig 5E), pathogenic mutations including both mild and severe mutational sites always demonstrate lower MSF values, except some pathogenic mutations at the Ca^2+^ binding domain. Some particular pathogenic mutations, located at minimal positions that correspond to hinge sites, have been found, such as M62, G63, S65, A377, D378, and H381 belong to severe mutational sites, and D60, V461, and G473 correspond to mild mutational sites. Structurally, these hinge mutational hotspots are located at the active site and the dimer interface. These conformational dynamic signature of disease-associated mutations have been revealed both at the genome[76] and proteome-levels.[31] Regarding the PRS analysis, two matrices were used to quantify the allosteric effect of each residue, which is effectiveness and sensitivity, for evaluating the propensity of residues to act as sensors or as effectors of allosteric signals. As shown in profiles in Fig 5F and G, the distribution of effectiveness and sensitivity for *ALPL* mutation hot spots shows different trends. In general, most of the pathogenic mutations match almost exactly with some peaks of effectiveness profile, with sharp dominant peaks being found for three severe mutational sites (G63, A377, and V459). On the other hand, the peaks of the sensitivity profile correspond to some control mutational sites, while pathogenic mutational sites have smaller sensitivity, such as V95 and G473 in the mild group and G63, S65, and V459 in the severe group. As two unusual ENM-based dynamics features, the distribution of mutations in MBS and stiffness profiles are shown in Fig 5 H and I. For the MBS profile, the overall distribution of pathogenic mutational sites is larger than that of control mutational sites, highlighting the importance of pathogenic mutational sites in maintaining the stability of the *ALPL* protein.

Lastly, the mapping of mutational sites onto stiffness demonstrates the relatively uniform distribution; thus, it is not easy to find hits to distinguish severe, mild, and control mutational sites by only considering mechanical stiffness.

In order to clearly observe the ability of dynamic parameters to distinguish mutations in different groups, statistical analysis was further performed on the data of severe, mild, and control mutational sites. As shown in Fig 6 (E), the control group had the highest mean MSF, with *P* = 1.8e-05 and 0.00021 by the Wilcoxon signed rank test, in comparison with mild and severe mutations, showing the significant differences between both pathogenic mutational sites and the control group. However, significant differences were not found between mild and severe mutational sites, with *P* =0.75. Similar significant results were also found for effectiveness (Fig 6F), sensitivity (Fig 6G), and MBS (Fig 6H), while significant differences in stiffness could not be found among the three groups of mutations (*P*> 0.01, Fig 6I).

In the present section, we have compared the conformational dynamics of the three types of *ALPL* mutations in terms of five ENM-based features. In general, the disease-related mutational sites are shared by lower conformational flexibility and propensity to act as sensors, but have high propensity to act as effectors and key nodes in maintaining stability. This suggests that the inclusion of ENM-based dynamics descriptors may increase the prediction of pathogenic mutations in *ALPL*.

### Machine Learning Analysis Determines Key Molecular Features for Classifying Different Mutation Types

In order to find the relationship between the above studied features, their redundancy and ability to classify and predict various mutation types, we first computed the pairwise correlations between different prediction scores by using Spearman’s rank correlation coefficient (Fig 7A). It is easy to comprehend that these features can be classified into three major types according to their related sequence, network, and dynamics information. Among sequence-based features, has high positive correlations with *MI* and RASA, but these three features all have negative correlation with conservation calculated by Consurf. For the four network parameters, ΔK, ΔB and ΔC show significant correlations between each other; only ΔCc is an independent factor. The dynamics-based matrices that show more complex correlations, have been divided into two groups: one includes stiffness, MSF, and sensitivity, and the other includes effectiveness and MBS. A highly positive correlation was predicted within each group, but a negative correlation was found between the two groups. The special index ΔΔG did not show any correlation with other features. Accordingly, this suggests that these features could complement our understanding of the molecular signatures of HPP-causing mutations.

**Fig 7.**
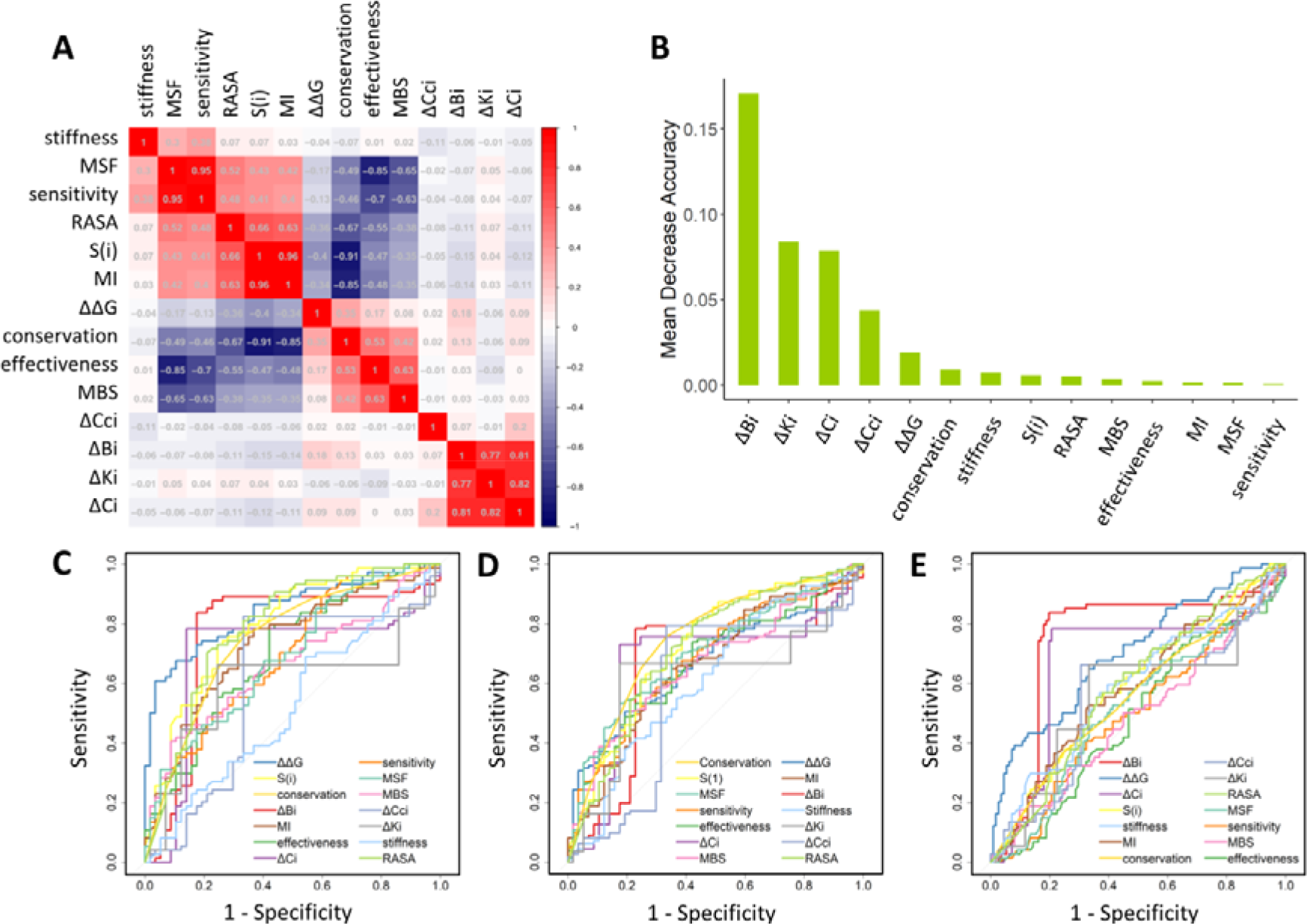
Performance evaluation of feature classification. (A) The heat map of pairwise Spearman’s rank correlation coefficients between different features. (B) Feature importance of all used features ranked based on Mean Decrease Accuracy in the RF classification. ROC curve AUCs for 14 features as a function of 1-specificity, including ROC curves for evaluating each feature in classifying *ALPL* mutations between (C) the control and severe groups, (D) control and mild groups, and (E) mild and severe groups.

Next, we employed a RF model to classify mild, severe, and control groups using the 14 features mentioned above (See S1 Text for detail method). Through the RFE algorithm, we considered classifying the model containing five variables, whose accuracy of repeated cross-validation was 80.5%, as the optimized one, which was used for comparison with the model using all features. To be more specific, the feature importance was ranked by the mean decrease accuracy (Fig 7B). Most interestingly, we found that the top four important features were all network-based features, including ΔK, ΔB, ΔC, and ΔCc, suggesting an essential role of protein network topology in controlling mutation events. ΔΔG is an important feature and is also an important determinant of protein stability in mutations. To our surprise, features relating to ENM protein dynamics were not ranked as important predictors. This may be caused by the loopy structure of the TNSALP protein. By considering only the top five features, the RF model yielded an accuracy of 96.8%. However, taking all features into account, the accuracy of the RF model dropped to 92.2%. The inclusion of more features reduces the predictive accuracy of the RF model, which led us to focus on the specific interpretability of each parameter.

Toward the interpretability of the classification effect of each parameter, we further evaluated the performance of the 14 features of the dataset composed of mild and severe mutations, mild and control mutations, severe and control mutations. To this end, the ROC curves with the AUCs of the 14 metrics for each comparison group were plotted, and the related values are listed in Table 1. First, we examined the difference in the AUCs for the 14 functional features of the severe and control groups (Fig 7C); ΔΔG showed appreciable performance with AUC 0.8447. Moreover, the *S*_(*i*)_, conservation score, ΔB also showed good performance with AUCs, all greater than 0.73. We then tested the performance of the 14 features between mild and control group mutations (Fig 7D), as well as the mild and severe group mutations (Fig 7E). In the classification of the mild group and control group, two sequence-based and four dynamics-based parameters showed moderate predictive performance. The AUC of the conservation score, entropy, MSF, effectiveness, sensitivity, and MBS were 0.7493, 0.7204, 0.7023, 0.6709, 0.6802, and 0.6666, respectively. By contrast, in the classification of the mild and severe groups, we found that ΔB showed the best performance, with an AUC = 0.7471, while, ΔΔG showed the second-best performance, with an AUC = 0.7048.

**Table 1.** The AUC of 14 characteristics or parameters in each comparison group. The characteristics that performed best in each group are highlighted in red.

| Features | Mild vs control | Severe vs control | Mild vs severe |
| --- | --- | --- | --- |
| conservation | **0.7493** | 0.7927 | 0.5632 |
| S(i) | 0.7204 | 0.7934 | 0.5906 |
| MI | 0.6633 | 0.7336 | 0.5841 |
| RASA | 0.5284 | 0.5043 | 0.5329 |
| $\Delta\Delta G$ | 0.6641 | **0.8447** | 0.7048 |
| $\Delta K_i$ | 0.6185 | 0.5991 | 0.5538 |
| $\Delta B_i$ | 0.6618 | 0.7636 | **0.7471** |
| $\Delta C_i$ | 0.6675 | 0.6961 | 0.6682 |
| $\Delta C_{c_i}$ | 0.5886 | 0.6081 | 0.5539 |
| MSF | 0.7023 | 0.6893 | 0.5140 |
| effectiveness | 0.6709 | 0.7009 | 0.4850 |
| sensitivity | 0.6802 | 0.6957 | 0.4968 |
| MBS | 0.6666 | 0.6632 | 0.4893 |
| stiffness | 0.619 | 0.5281 | 0.5860 |

**Table 2.**
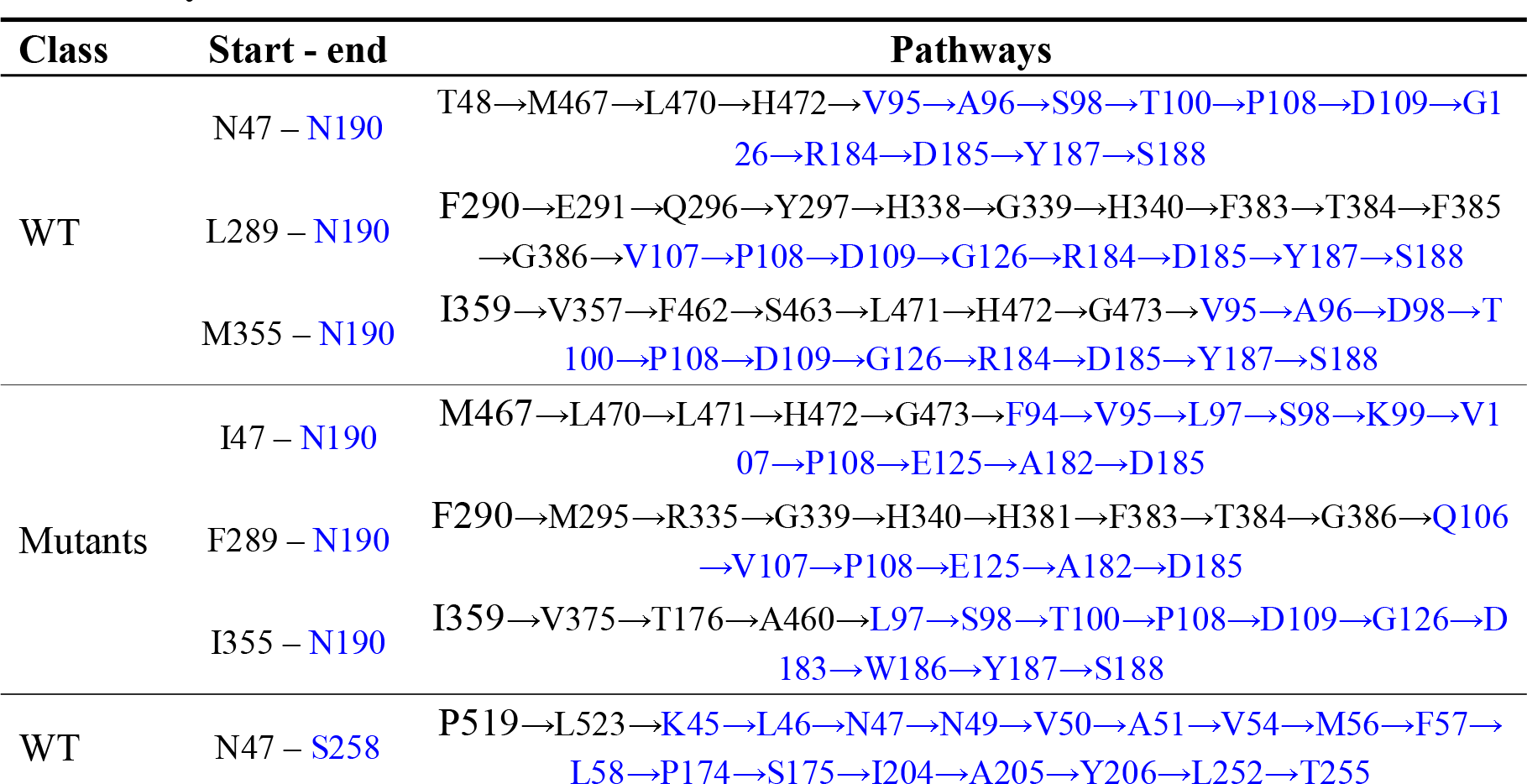
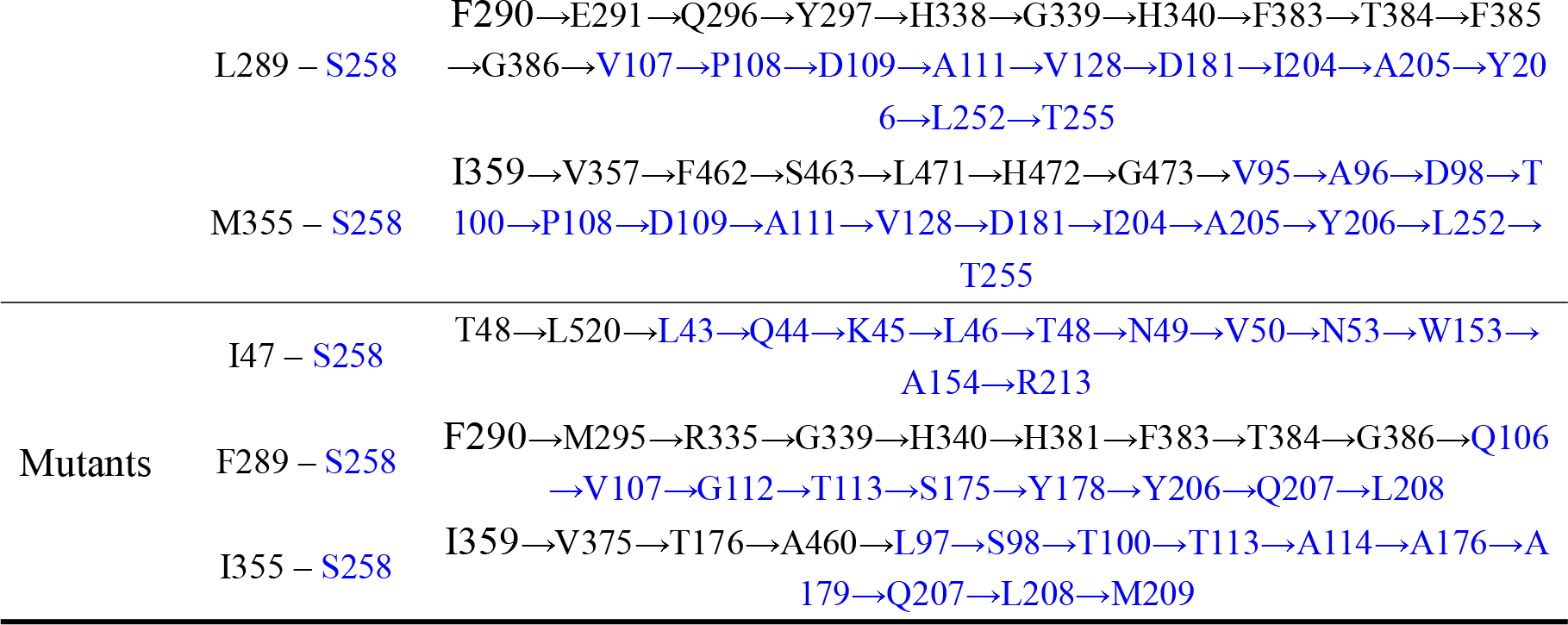
The constituent residues of the shortest pathway from the three severe mutational sites (N47, L289, and M355 to N190/S258). Residues in the two chains are denoted by different colors.

In the features analysis, we argue that protein sequence, structure and dynamics all play important roles in the understanding of *ALPL* mutations and their resultant phenotypes. In our study of ALPL mutations, the RF model only included network topological and ΔΔG features that can predict the phenotypic severity of hypophosphatase. The most interesting finding is that ΔΔG is a good indicator for identifying disease mutations, while ΔB is the best indicator for classifying mild and severe mutations.

### Atomistic Simulations and Network Modeling Determine the Effects of Severe Phenotype-Associated Mutations on Allosteric Communications

Based on previous results, some severe phenotype-related mutations have relatively low ΔΔG but higher ΔB values in network topology. To gain energetic and topological insights, we have compared ΔB and ΔΔG values for all mutations. The scatterplot revealed that N47I, L289F and M355I were three severe mutations with large ΔB but low ΔΔG (Fig 8A). In order to determine the dynamics effects induced by severe phenotype-associated mutations, we performed a set of four independent, all-atom MD simulations for WT and three mutant TNSALPs caused by N47I, L289F, and M355I variants (Fig 8). The RMSD values of Cα atoms served as an overall measurement of the departure of the structures from the initial coordinates (S5A Fig). All the systems became for a duration of 30 -50 ns, and the RMSD values became stable during the last 50 ns of the simulations. The conformational dynamics results (S5B Fig) revealed that the profile of RMSF is very consistent in the WT and mutant systems, and no conformational change occurred at the active site, further explaining why ΔΔG for these severe mutations were small. The differential fluctuation ΔRMSF, with respect to WT, showed considerable changes in regions around the ion binding pocket and the Ca^2+^ binding site located in the other monomer, with two of the largest peaks at residues, N190 and S258 (Fig 8B). This suggests the existence of allosteric communication between the mutant sites and these two regions. Accordingly, in the four systems, we selected mutant sites in the chain A as starting points and N190 and S258 of the chain B as the target residues to further identify specific allosteric pathways caused by severe phenotype-associated mutations.

**Fig 8.**
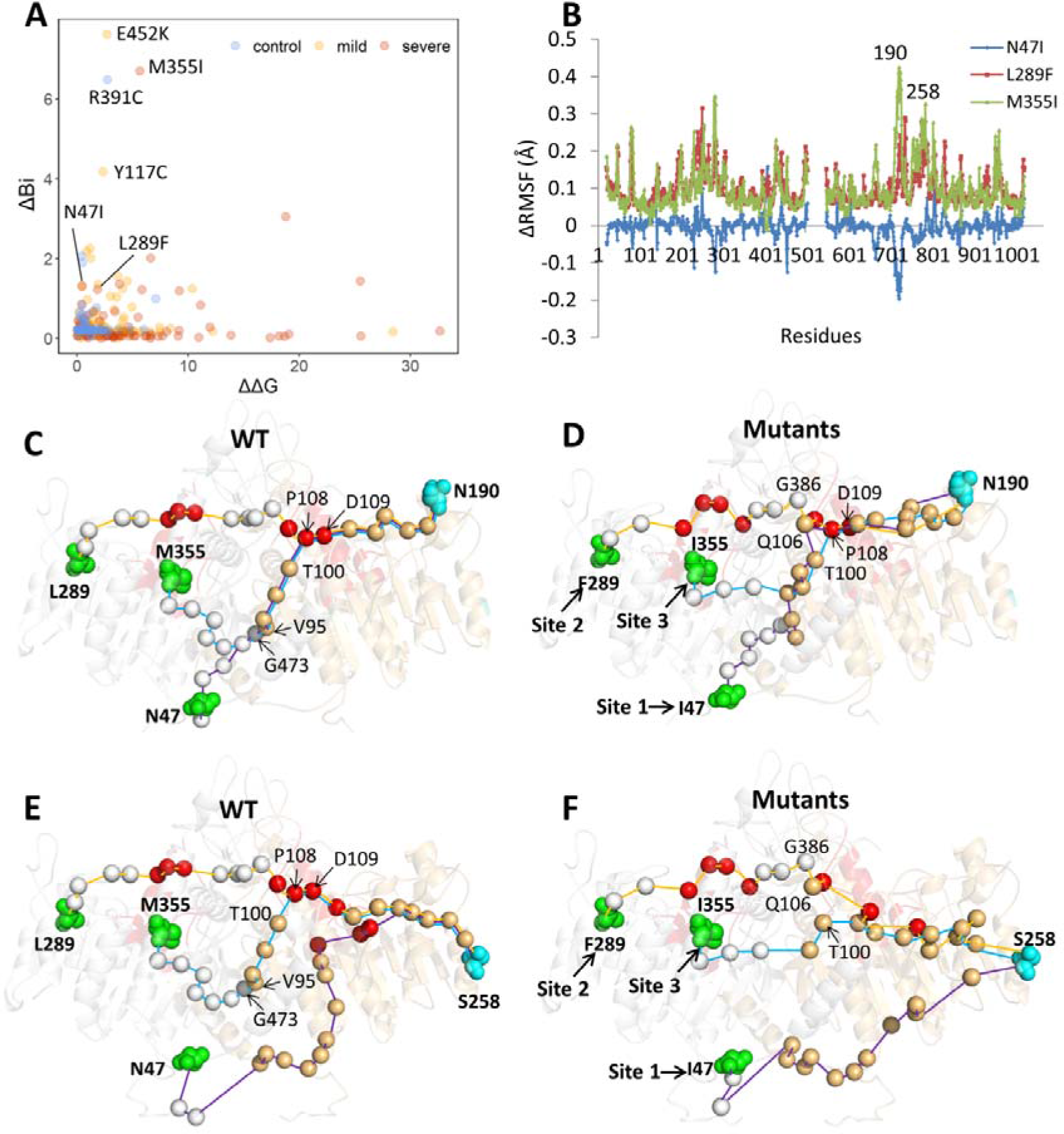
(A) The distribution of ΔΔG and ΔB for *ALPL* mutations. The scatterplot showing distribution of ΔB vs ΔΔG of different mutation types. Severe, mild and control mutations are depicted in red, yellow, and blue, respectively. N47I, L289F, and M355I are three severe mutations with low ΔΔG and high ΔB. Among the three significant mutations predicted by the scatterplot, two (E452K and R391K) were not originally included in the severe mutation group but were validated as two severe mutations in the newly collected clinical samples. (B) Differential RMSF (ΔRMSF) of N47I (blue), L289F (red), and M355I (green), with respect to 36 WT. Allosteric paths originating at three mutational sites and terminating at N190 in the WT (C) and mutant states (D), as well as terminating at S258 in the WT (E) and mutant states (F), respectively. The TNSALP structure is depicted as represented by semi-transparent colored cartoon, and the starting and ending residues of all the paths are represented as green and cyan spheres, respectively. The alpha-carbon of the path through the residues is shown as silver (chain A) and orange (chain B) spheres, in which active sites are represented by red spheres.

The shortest path algorithm based on dynamic network models was employed to identify specific allosteric pathways. As shown in Fig 8 (C-F), the inter-molecular interaction of A: G473-B: V95 was predicted as the key bridge in WT, and the other interaction of A: G386-B: Q106 as the key bridge in mutants. It was, therefore, determined that WT and mutant TNSALPs use different allosteric interfaces for signal transmission. Overall, pathway plasticity was found to exist, although the structures between WT and mutants were well conserved, conforming to the recent function-centric allosteric regulation study.[77] Among these pathways, a common region including three active site residues (T100→P108→D109) corresponding to mutational sites was also found, suggesting that these mutational paths may lead to the same functional state. In particular, the molecular signatures showed that T100, P108, and D109 have relatively high effectiveness and ΔB, demonstrating that these nodes are key sites for structural signal propagation.

The comparison of allosteric pathways reveals several important insights. First, by using the same starting mutational sites and ending points, the mutant TNSALPs exhibit shorter allosteric pathways, which is in accordance with our hypothesis that the mutants caused by severe mutations have higher allosteric propensities. The system with the largest reduction in signaling path nodes is the mutant caused by M355I, with has the highest ΔB and a relatively small ΔΔG. In WT, the path from M355 to S258 passed through the nodes F462→S463→L471→H472→G473 on the dimeric interface of the chain with mutations and the active sites P108 and D109 in the other chain (Fig 8E). In contrast, none of these sites participated in the signaling paths in the mutants. Hence the whole length becomes shorter and straight (Fig 8F). The same situation occurs in the path from M355I to N190 (Fig 8C, D). Second, the pathways in mutants involve fewer active sites, interfacial residues, and mutational sites. This comparison further highlights these functional regions as hubs for long-range allosteric communication in WTs, while it also means that the functional role of these reduced nodes may be lost if a severe mutation occurs. The extreme situation is observed for N47I, in which the path from I47 to S258 does not pass through the active area and employs the shortest and most straight path. Lastly, the path patterns are more robust in WT than in mutant states, suggesting that severe mutations introduce more pathway plasticity. For example, the shortest pathways from residues N47, L289, and M355 to N190 of the other chain in the WT shared some similar nodes, including P108, D109, G126, R184, D185, Y187, and S188, in the other chain, with most of located at the active site or ion binding pocket, and with all crossing the same area and to the ending residue N190 (orange circle region in Fig 8C and D). However, the situation was different in mutants. Fewer residues were shared in the three mutants, only residue P108 showed in the shortest pathway from I47, F289, and I355 to N190. The situation was similar in the paths from residues, I47, F289, and I355 to S258 of the other chain. Three paths shared residues, I204, A205, Y206, L252, and T255 of the other chain in the WT and none in the mutants.

Taken together, the results suggeted that severe *ALPL* mutaions may affect the pathway of signal transduction between the two monomers of TNSALP. Allosteric pathways in WTs are robust and involve some functionl sites, consisting of active, interfacial, and mutational residues. In the allosteric mutant states, the overall pathway pattern is more “flexible” with the shorter pathways involving fewer functional sites, resulting in the severe phenotype.

### Assessment of Allosteric Effects of Severe Mutational Sites

As shown in Fig 8 (A), among the four mutations (E451K, M355I, R391C andY117C) with largest ΔB but low ΔΔG values, only M355I related to severe HPP phenotype. At the time of waiting, the ALPL mutation database has been updated by including several more severe mutations. We were surprised to find that the new patients with severe phenotype related to E452K and R391C, were reported as mild and control mutations previously. To further investigate their allosteric effects and whether is an intrinsic dynamic of these severe mutational sites, the single-residue perturbation analysis was performed by using the AlloSigMA. The results showed that perturbations of all these five mutational sites induced inter molecular allosteric effects (Fig 9) in the form of both positive and negative modulation with a mean value of ΔG (N47→chain B) = 0.89 kcal/mol, ΔG (N289→chain B) = 0.11 kcal/mol, ΔG (N335→chain B) = 0.37 kcal/mol, ΔG (R391→chain B) = 0.13 kcal/mol and ΔG (E452→chain B) = 0.08 kcal/mol, respectively. The analysis exhibited an evident influence on the stability of some functional regions, including the Ca^2+^ binding and crown domain, thereby revealing that severe mutations can induce changes in the stability of other sites, and affect the catalytic activity of proteins.

**Fig 9.**
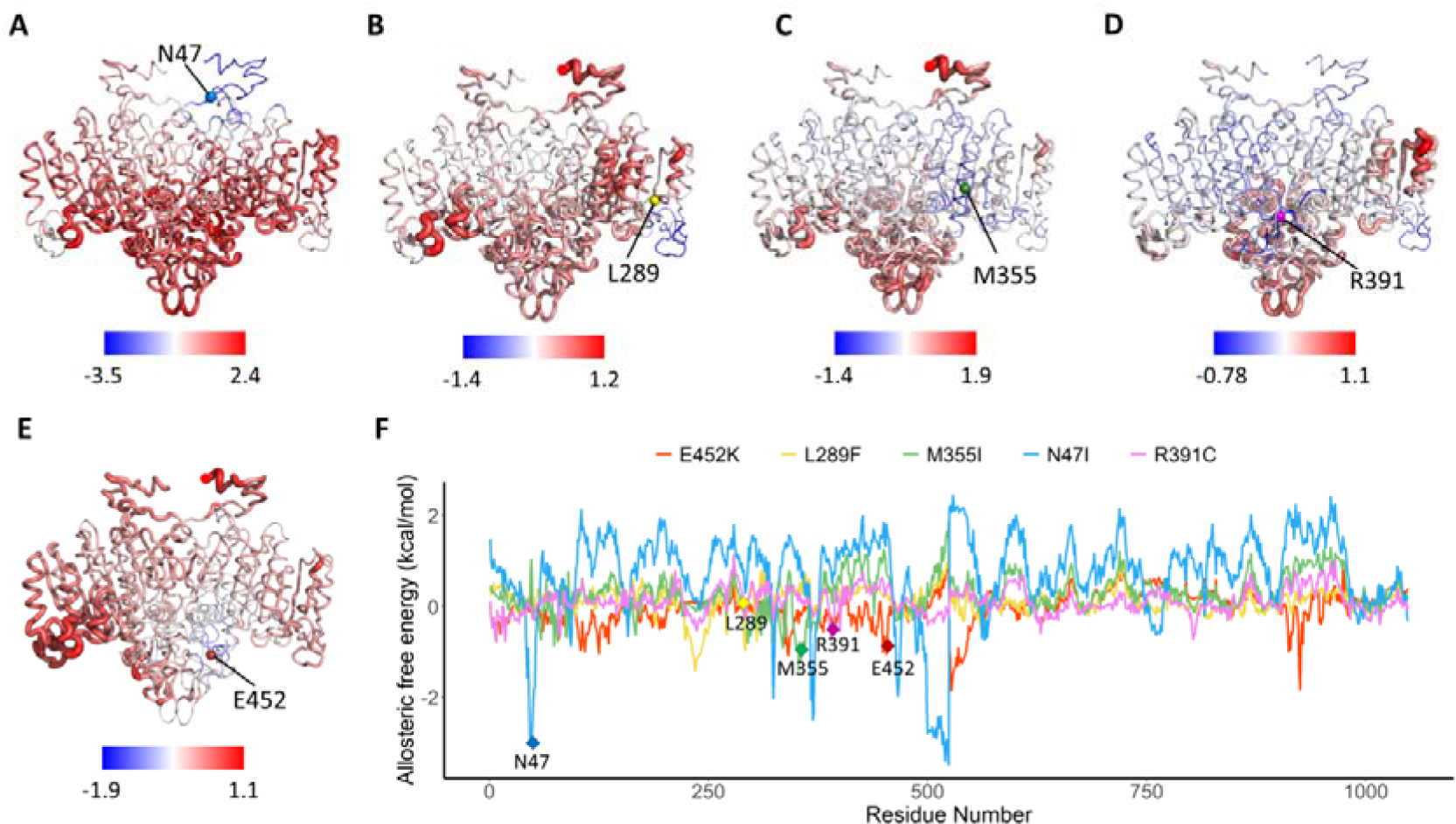
Allosteric effects of three studied severe mutations (N47I, L289F and M355I) and two predicted severe mutations (R391C and E452K) calculated by AlloSigMA. Cartoon structures of the TNSALP protein colored according to their free energy values obtained for the cases of (A) N47I, (B) L289F, (C) M355I, (D) R391C and (E) E452K, while blue color indicates negative allosteric free energy and red color indicates positive modulation. (F) Their free energy profiles are illustrated graphically with residue index (chain A: 1-524; chain B: 525-1048) on the x-axis and Δg value on the y-axis. Blue, yellow, green, pink and red profiles represent the result for N47I, L289F, M355I, R391C and E452K respectively.

These findings once again suggest that ΔB is a good and facile indictor for predicting severe mutations, whose molecular pathogenicity does not alter the local stability at active sites so as to change the protein folding energy, but instead generates alternative and long-range molecular effects.

## Conclusions

As the biological hallmark of HPP, the reduced activity of TNSALP is caused by loss-of-function mutations in the *ALPL* gene; varying levels of reduced activity are related to mild or severe HPP phenotypes. Despite some progress, more reseach is needed for comprehensive understanding of genotype-phenotype interrelationship in HPP. Thus, in this study, the functional landscape of *ALPL* mutations was established by investigating a set of features generated from the protein sequence, network topology, and ENM dynamics calculation, and their relationship with different HPP phenotypes. Based on these features, a machine learning classifier was developed not only to examine the relationship between the pathogenicity status of mutations and their biophysical attributes, but also for the prediction of mutations in the control, mild, and severe groups. We have found that in addition to ΔΔG as the commonly used predictor, co-evolutionary conservation and network-based features also yield strong signals in machine learning predictions and provide orthogonal information.

Further structure-function studies including MD simulation and structural-communication pathway analysis of mutant TNSALPs, have supported our arguments that mutational hotspot sites often correspond to global mediators of allosteric interactions. These findings suggest that the study of molecular signatures and allosteric regulation of *ALPL* mutations may be a step toward defining a greater quantitative genotype-phenotype interrelationship in HPP.

Through the large-scale analysis of disease-causing *ALPL* mutations, we propose the following possible molecular principles underlying HPP-related mutations. First, HPP pathogenicity is largely due to the structural instability of the TNSALP caused by *ALPL* mutations, which have variable effects on enzyme activity. Thus, for such cases, ΔΔG is a satisfactory and acceptable index for predicting pathogenic mutations, especially for distinguishing mutations between the control and severe groups. There is also the possibility of extrapolating our methods of using ΔΔG to estimate change in protein stability upon mutations as a popular way to predict pathogenic mutations in other diseases. Second, the most interesting finding in our opinion is that ΔB has been predicted as a good network indicator to distinguish mild and severe groups in pathogenic mutations. We speculate that mutations in the severe group have stronger allosteric effects than mutations related to the milder forms of HPP, severing as “allosteric mutations”.[78, 79] Analysis of allosteric properties of severe mutations adds an additional confirmatory layer to segregate variants in “mild” and “severe” states, thereby furthering understanding of the allosteric basis of loss-of-function mutations. Third, for the classification of mild mutations and differentiation from the control group, co-evolutionary conservation has been shown to be the most important predictor, thereby suggesting that mild pathogenicity may be related to amino acid changes with small evolutionary substitution probability. However, to reach the elusive goal of establishing the precise relationship between *ALPL* mutation genotypes and HPP phenotypes and a more reliable prediction model or score, like as protein regulatory and functional binding sites prediction done,[80] more clinical data on mutations[81] and data on enzyme activity[13] are needed.

## Supporting Information

S1 Table. Collected data set of ALPL mutations.

S1 Fig. Distribution of ALPL mutations in terms of WT TNSALP sequence and structure.

S2 Fig. Evolutionary conservation analysis performed for TNSALP protein P60484 with 403 aa using ConSurf.

S3 Fig. Ramachandran plot of TNSALP.

S4 Fig. Comparisons of different molecular signatures for ALPL mutations.

S5 Fig. Conformational dynamics of TNSALPs.

## Acknowledgments

This work was supported by the National Natural Science Foundation of China (31872723), a Project Funded by the Priority Academic Program Development (PAPD) of Jiangsu Higher Education Institutions, the China Postdoctoral Science Foundation (2016M590495), and the Jiangsu Planned Projects for Postdoctoral Research Funds (1601168C).

## Author Contributions

The manuscript was written through contributions of all authors. All authors have given approval to the final version of the manuscript. ‡These authors contributed equally.

## Notes

The authors declare no competing financial interest.

## Abbreviations

HPP, Hypophosphatasia; TNSALP, tissue-non-specific alkaline phosphatase; MSA, Multiple Sequence Alignment; SASA, solvent-accessible surface area; PSN, protein structure network; ENM, elastic network model; RF, Random Forest; ROC, Receiver Operating Characteristics; AUC, Area Under Curve.

## Notes

### Competing Interest Statement

The authors have declared no competing interest.

